# An improved genome assembly of the saguaro cactus (*Carnegiea gigantea* (Engelm.) Britton & Rose)

**DOI:** 10.1101/2023.04.11.536419

**Authors:** Dario Copetti, Alberto Búrquez, Kentaro K. Shimizu, Rod A. Wing, Michael J. Sanderson, Martin F. Wojciechowski

**Affiliations:** Arizona Genomics Institute, School of Plant Sciences, University of Arizona, Tucson, AZ 85721, USA; Department of Evolutionary Biology and Environmental Studies, University of Zurich, 8057 Zürich, Switzerland; Instituto de Ecología, Unidad Hermosillo, Universidad Nacional Autónoma de México, Hermosillo, Sonora, Mexico; Center for Desert Agriculture, Biological and Environmental Sciences & Engineering Division (BESE), King Abdullah University of Science and Technology (KAUST), Thuwal, 23955-6900, Saudi Arabia; Department of Ecology and Evolutionary Biology, University of Arizona, Tucson, AZ 85721, USA; School of Life Sciences, Arizona State University, Tempe, AZ 85287, USA

**Keywords:** Cacti, genome assembly, saguaro cactus, Cactaceae, *Carnegiea gigantea*

## Abstract

We present an improved genome assembly of the saguaro cactus (*Carnegiea gigantea* (Engelm.) Britton & Rose), obtained by incorporating long-read PacBio data to the existing short reads. The assembly improves in terms of total size, contiguity, and accuracy, allowing to extend the range of sequence analyses beyond the single-gene scale. Consequently, the assembly is 16% larger and has 20% more genes, expanding the resources for a neglected yet very remarkable plant family such as Cactaceae.

**Species taxonomy:** Eukaryota; Viridiplantae; Streptophyta; Streptophytina; Embryophyta; Tracheophyta; Euphyllophyta; Spermatophyta; Magnoliopsida; Mesangiospermae; eudicotyledons; Gunneridae; Pentapetalae; Caryophyllales; Cactineae; Cactaceae; Cactoideae; Echinocereeae; Carnegiea gigantea (Engelm.) Britton & Rose) (also known as saguaro cactus) (NCBI txid: 171969).

## Background

Technological improvement in DNA sequencing methods has allowed the large-scale development of genomic resources for non-model organisms, secondary crops, and ultimately any species. However, limited financial availability and difficulty in sample preparation still prevent the genomes of certain organisms from being fully characterized. Despite being an extreme example of physiological and morphological adaptation, until recently little or no genomic resources were available for cactus species^1–4^.

In this short report, we describe the improvement of the genome assembly of *Carnegiea gigantea* (Engelm.) Britton & Rose – the saguaro cactus. SGP5, the specimen that was sequenced in the initial assembly^1^ (Figure 1) succumbed to a thunderstorm in August 2016 and attempts in clonally propagating the specimen failed. However, we were able to germinate a few seeds from an SGP5 fruit and one single seedling (named SGP5p, with p for progeny) was sacrificed with the goal of producing long-read data to improve the assembly of the parental genome. Aware of the presence of paternal variants in SGP5p’s long read data, we performed error correction with SGP5’s short reads to restore the set of SGP5’s variants on a longer DNA sequence backbone. Upon validating the assembly, we performed de novo annotation of the genome. Though still partially complete, this resource represents the highest quality public genome assembly for the whole Cactaceae family. With the hope that soon also cacti will have a worthy reference assembly to serve as an anchor for research programs, we release to the public an updated genome assembly and annotation of the iconic saguaro cactus.

**Figure 1.**
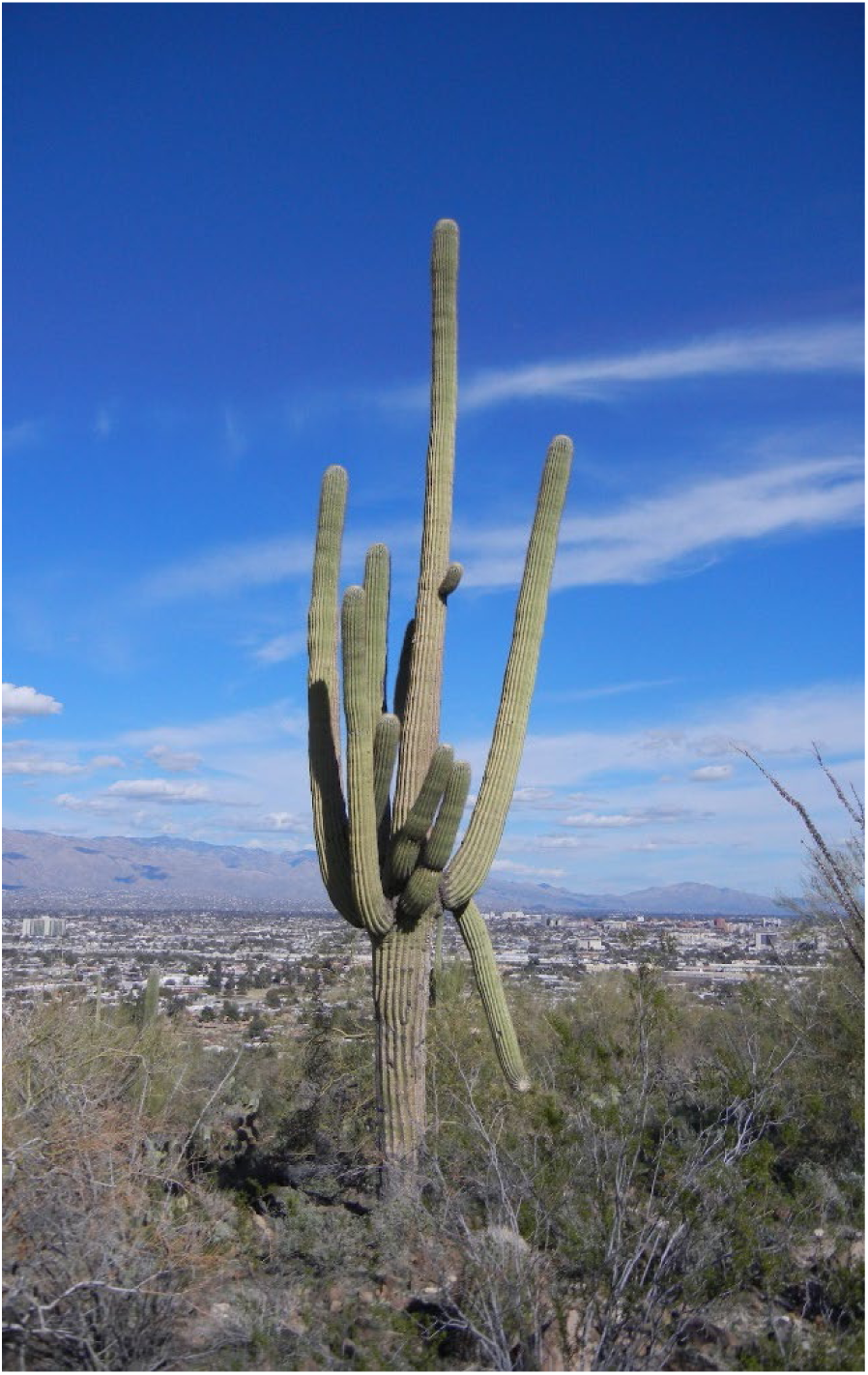
The SGP5 individual on Tumamoc Hill, Tucson, Arizona. Photo: MF Wojciechowski.

## Genome sequencing report

DNA from a progeny (SGP5p) of SGP5 was sequenced with the CLR technology on a PacBio Sequel system. The long-read dataset consisted of 1.6 million reads, spanning 26.8 Gb (about 19 genome equivalents), with a N50 and a median read length of 25.5 and 13.6 kb, respectively. The long reads were combined with the PE data from^1^ in a hybrid assembly (named SGP5p v2), consisting of 5,942 scaffolds spanning 1.140 Gb – 81% of the estimated saguaro genome size. Half of the assembly was included in the 649 longest sequences (the shortest of which was spanning 467 kb), and 90% of the bases were included in the 2,695 sequences (the shortest was 92 kb in size, Table 1). In comparison with the v1.3 assembly^1^ (obtained using only short reads), this updated assembly contained 16% more bases and was a significant improvement in contiguity, having an almost 10-fold reduction in number of sequences and a 7 to 11-fold improvement in the Nx metrics (Table 1). The new assembly also showed a strong reduction (151-fold reduction) in the number and span of sequence gaps.

**Table 1.** Metrics of the two saguaro assemblies and of the dragon fruit assembly after splitting the sequences at large gaps.

|  | *C. gigantea* |  | *S. undatus* |
| --- | --- | --- | --- |
|  | SGP5 v1.3 | SGP5p v2 | D. Bowie |
| **Number of contigs** | 57,405 | 5,942 | 88,634 |
| **Total length (Mb)** | 980.3 | 1,140.7 | 1,164.3 |
| **Longest sequence (bp)** | 648,566 | 5,085,408 | 654,357 |
| **Mean sequence (bp)** | 17,078 | 191,979 | 13,136 |
| **GC (%)** | 36.8 | 37.1 | 37.4 |
| **N50** | 61,549 | 467,946 | 46,368 |
| **L50** | 4,574 | 649 | 6,532 |
| **N70** | 34,419 | 265,749 | 20,637 |
| **L70** | 8,804 | 1,291 | 13,980 |
| **N90** | 8,017 | 92,355 | 5,082 |
| **L90** | 19,447 | 2,695 | 37,439 |
| **Gaps** | 60,007 | 396 | 16,628 |
| **N bases** | 18,951,006 | 39,703 | 166,280 |
| **% of N bases** | 1.9 | 0.003 | 1.4 |
| **BUSCO complete** | 92.9 | 95.0 | 96.9 |
| **single copy** | 89.4 | 89.7 | 92.1 |
| **duplicated** | 3.5 | 5.3 | 4.8 |
| **fragmented** | 4.8 | 1.5 | 2 |
| **missing** | 2.3 | 3.5 | 1.1 |
| **Gene models** | 28,292 | 34,209 | 28,992 |

Besides saguaro, the only publicly available highly contiguous genome assembly of a cactus species is *Selenicereus undatus* (Haw.) D.R. Hunt, developed with 10X Genomics and Dovetail Chicago and Hi-C data^3^. Other assemblies, published in^1,4^ are extremely fragmented and incomplete, barely represent the genic portion, and will not be considered here as a comprehensive representation of a plant genome. The dragon fruit assembly manuscript lacks substantial details in the how contigs and scaffolds were obtained. Together with the large number of gaps, this work warrants caution when utilizing such resource for analyses beyond the gene-level scale. To offset for the spatial organization revealed by Hi-C scaffolding and have a more appropriate comparison of the assigned bases in the *S. undatus* assembly when comparing it to SGP5p v2, we split the dragon fruit scaffolds at gaps of 100 bp or more. This removed 162 Mb (12% of the total assembly size) of unassigned bases. The resulting “contig-level” assembly contained more contigs than the first saguaro assembly released, for a total assembly size comparable to SGP5p v2, and had contiguity values 10-fold lower than SGP5p v2 (Table 1). The cumulative length plot (Figure 2) clearly shows the higher contiguity of SGP5p v2 compared to both its predecessor and *S. undatus* assemblies.

**Figure 2.**
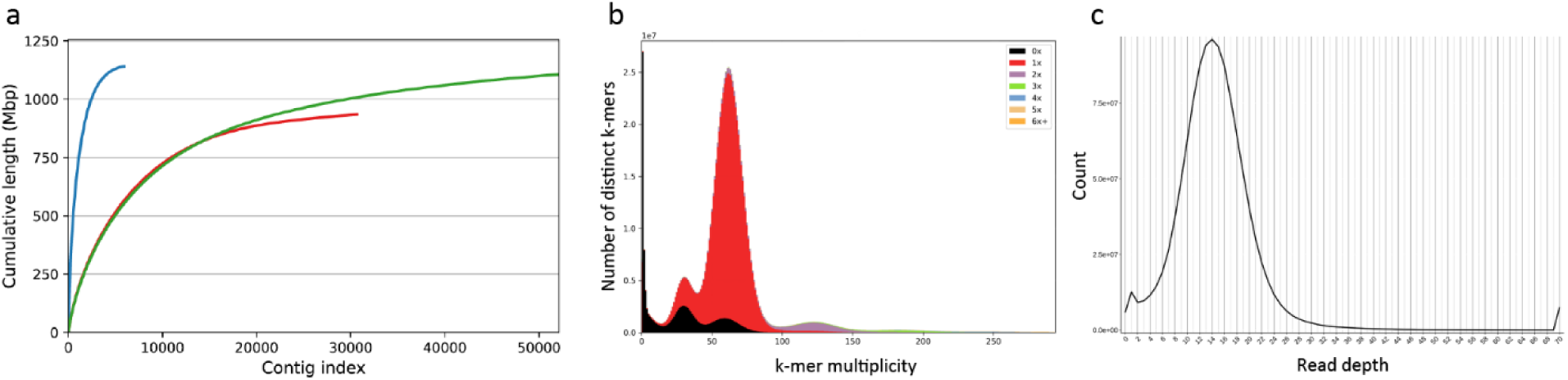
a) QUAST cumulative plot of the three most complete cactus genome assemblies. SGP5 v1.31 profile is in red, SGP5p v2 (current study) is in blue, and the dragon fruit contig-level assembly is in green. Panel b shows the KAT coverage plot of SGP5p v2.

Compared to the first saguaro assembly, the SGP5p v2 assembly had a higher BUSCO score, derived from less fragmented and more duplicated single-copy ortholog models (Table 1). Assembly completeness was estimated by comparing 21-mers present in the error-corrected short reads and in the final assembly, resulting in 94% completeness – an acceptable value assuming the assembly contained only one of the allelic variants in the haploid assembly. As expected, the assembly retained about half of heterozygous k-mers (47% missing, 53% present, peak at x = 30 in Figure 2 b), while less than 4.7% of the homozygous k-mers were missing and 2.3% were duplicated (black and purple area at x = 61, respectively). A majority fraction (68%) of the duplicated k-mers was present twice in the assembly (purple area at x = 120) – supporting a good representation of the repeated sequences in the assembly.

To assess for inadvertent retention of sequences of both allelic variants at heterozygous genomic regions, long reads were aligned to the assembly. The coverage distribution plot of the alignment data showed the absence of regions that have a coverage value half the one in the haploid assembly (Figure 2 c). Evidence from both the k-mer and the coverage analyses together support the claim that the SGP5p v2 assembly represents a haploid set of the saguaro genome.

Gene annotation identified 34,209 protein-coding gene models in the SGP5p v2 assembly. The quality of the predictions is supported by 92.75% of the models with annotation edit distance at or below 0.5, and 18.92% at or below 0.2. The average and median exon size were 249 and 351 bp, respectively. Exons span 46.6 Mb. Compared to the previous version, this updated assembly contained 20% more gene models (28,292 in SGP5 v1.3) that span a wider proportion of the assembly (4.1% vs 3.8%).

The amount of sequence represented by transposable elements and repeated sequences was estimated both with homology and with structure-based methods. Overall, 63% of the assembly was composed interspersed repeats, of which the dominant fraction (68%) was LTR-retrotransposons, followed by DNA transposons (20%). Helitron and repeated sequences occupied about 4% of the assembly each, while non-LTR retroelements were detected in less than a percentage point (Table 2).

**Table 2.** Diversity and abundance of different transposable elements and repeat classes in the SGP5p v2 assembly.

| Class |  | Repeat count | Masked bases (Mb) | Assembly % masked |
| --- | --- | --- | --- | --- |
| LTR | Copia | 86,692 | 116.7 | 10.2% |
|  | Gypsy | 183,698 | 224.9 | 19.7% |
|  | unknown | 263,034 | 148.2 | 13.0% |
| TIR | CACTA | 82,830 | 34.8 | 3.1% |
|  | Mutator | 118,441 | 52.9 | 4.6% |
|  | PIF_Harbinger | 68,566 | 25.4 | 2.2% |
|  | Tc1_Mariner | 21,822 | 7.7 | 0.7% |
|  | hAT | 48,413 | 20.7 | 1.8% |
| nonLTR | LINE_element | 1,352 | 0.7 | 0.1% |
|  | unknown | 1,363 | 1.4 | 0.1% |
| nonTIR | Helitron | 92,358 | 49.0 | 4.3% |
| Repeat region |  | 128,757 | 38.6 | 3.4% |
| Total |  | 1,097,326 | 721.0 | 63.2% |

## Methods

### Sample acquisition and nuclei acid extraction

Seeds from the SGP5 plant were germinated and grown in a greenhouse. A 2.5-year-old, ∼10 cm tall seedling was selected for DNA extraction using the Qiagen® Genomic-tip extraction kit. The specimen was named SGP5p.

### Sequencing

A PacBio WGS library was constructed with SGP5p genomic DNA and sequenced with the Sequel® binding kit v3.0 in three SMRT cells 1M for 10 h on a PacBio Sequel instrument at Oregon State University, USA.

### Genome assembly

The genome was assembled with MaSuRCA^5^ (v.3.3.9) with NUM_THREADS = 40 JF_SIZE = 14000000000. Input data were the Illumina data from^1^ (SRR5036292, SRR5036295, and SRR5036296) and the newly-generated PacBio data (SRR17493743, SRR17493742, and SRR17493741). For the coverage analysis, MaSuRCA error-corrected megareads were aligned to the assembly with minimap2^6^ (v.2.17-r941), alignments were parsed with SAMtools^7^ (v.1.13, subcommands view, sort, index) and bedtools^8^ (v. 2.30.0, subcommand coverage). Mean coverage in 10 kb regions was plotted. Chloroplast and mitochondria genomes were downloaded from^9^ and NCBI accession FP885845 (*Beta vulgaris*), respectively, and used as a database to identify organellar sequences in the SGP5p v2 assembly via BLASTN^10^ (v.2.11.0+). One scaffold contained the whole chloroplast genome, and the sequence was edited to reflect the same start position as in other chloroplast genomes. Assembly errors were corrected by four rounds of subread alignment (with minimap2) and consensus calling with Racon^11^ (v.1.4.20). The resulting sequence was then incorporated together with the nuclear genome assembly. BLASTN (-evalue 1e-5) alignment revealed a single scaffold with similarity to the mitochondria genome.

Properties of the assembly were assessed with BUSCO^12^ (v.5.2.2, --augustus_species tomato –l embryophyta_odb10 -m genome -e 1e-10) and KAT^13^ (v2.4.2, subcommand comp -H 10000000000 -h -m 21). Assembly comparison was performed with QUAST^14^ (v. 5.0.2 --large --conserved-genes-finding --no-snps --no-sv --no-read-stats).

### Genome annotation

Protein-coding regions were predicted with MAKER (v2.31.10). RNA-Seq data from^1^ (SRR5134694, SRR5134696, SRR5134695, and SRR5134693) were aligned to the assembly with Hisat2^15^ (v2.2.1, -dta --max-intronlen 100000), then parsed with SAMtools (subcommands view, sort, index), Stringtie^16^ (v2.1.7, -m 150), and Gffread^17^ (v0.12.7) to obtain hints for gene prediction. Other sets of evidence were: gene models, *Lophophora williamsii* unigenes and a plant protein library, Augustus and SNAP ab initio predictions from^1^. The assembly as masked for repeats with RepeatMasker (https://www.repeatmasker.org/, v. open-4.0.7, Search Engine: NCBI/RMBLAST [2.6.0+]) and the repeat library from^1^. MAKER gene models that did not start with a methionine were removed, as well as models with similarity to TE genes for more than 40% BLASTP (-evalue 1e-5) identity, of at least 33 aa, and spanning at least 40% of the query length.

A global TE annotation was performed with EDTA^18^ (v.2.0.1, --cds [CDS from gene annotation] --anno 1 -- evaluate 1 --sensitive 1), non-coding RNAs were predicted with Infernal^19^ (v.1.1.4, Rfam database v14.6, as cmscan -Z 10000 --cut_ga --rfam --nohmmonly --fmt 2 --oskip) and tRNA-Scan-SE^20^ (v.2.0.9).

## Data availability

This Whole Genome Shotgun project has been deposited at DDBJ/ENA/GenBank under the accession JAKOGI000000000. The version described in this paper is version JAKOGI010000000. The assembly and annotation are available at https://doi.org/10.5061/dryad.hdr7sqvnp.

## Acknowledgements

The authors are grateful to the Center for Genome Research and Biocomputing at Oregon State University for long-read data generation and to the IT Support Group D-HEST of ETH Zurich for computational support. This work was funded by the NSF EAGER award # 1735604 to MFW and MJS.

